# Complete avian malaria parasite genomes reveal features associated with lineage specific evolution in birds and mammals

**DOI:** 10.1101/086504

**Authors:** Ulrike Böhme, Thomas D. Otto, James Cotton, Sascha Steinbiss, Mandy Sanders, Samuel O. Oyola, Antoine Nicot, Sylvain Gandon, Kailash P. Patra, Colin Herd, Ellen Bushell, Katarzyna K. Modrzynska, Oliver Billker, Joseph M. Vinetz, Ana Rivero, Chris I. Newbold, Matthew Berriman

## Abstract

Avian malaria parasites are prevalent around the world, and infect a wide diversity of bird species. Here we report the sequencing and analysis of high quality draft genome sequences for two avian malaria species, *Plasmodium relictum* and *Plasmodium gallinaceum*. We identify 50 genes that are specific to avian malaria, located in an otherwise conserved core of the genome that shares gene synteny with all other sequenced malaria genomes. Phylogenetic analysis suggests that the avian malaria species form an outgroup to the mammalian *Plasmodium* species and using amino acid divergence between species, we estimate the avian and mammalian-infective lineages diverged in the order of 10 million years ago. Consistent with their phylogenetic position, we identify orthologs of genes that had previously appeared to be restricted to the clades of parasites containing *P. falciparum* and *P. vivax* - the species with the greatest impact on human health. From these orthologs, we explore differential diversifying selection across the genus and show that the avian lineage is remarkable in the extent to which invasion related genes are evolving. The subtelomeres of the *P. relictum* and *P. gallinaceum* genomes contain several novel gene families, including an expanded *surf* multigene family. We also identify an expansion of reticulocyte binding protein homologs in *P. relictum* and within these proteins, we detect distinct regions that are specific to non-human primate, humans, rodent and avian hosts. For the first time in the *Plasmodium* lineage we find evidence of transposable elements, including several hundred fragments of LTR-retrotransposons in both species and an apparently complete LTR-retrotransposon in the genome of *P. gallinaceum*.

## Introduction

Malaria parasites of birds are more widespread, prevalent and genetically diverse than those infecting other vertebrates (Bensch et al. 2009); they are present in all continents except Antarctica, and in some populations up to 98% of birds within a species may be infected (Glaizot et al. 2012). However, there is considerable variation in their distribution across different host species. *Plasmodium relictum*, for example, infects a broad range of avian species, it has been found in birds of 11 orders, e.g. Passeriformes (Bensch et al. 2009) but *P. gallinaceum* has only been found in four species, including wild jungle fowl of Southern Asia and domestic chickens (Springer 1996).

The first avian malaria parasites were discovered in the late 19th century, shortly after the discovery of human malaria parasites. In the early 1900’s avian malaria became a prominent experimental model to study malaria biology (Huff and Bloom 1935; Raffaela and Marchiafava 1944) as well as for the routine testing and development of the first antimalarial drugs (Marshall 1942). Avian malaria is also a unique model to understand the ecology and evolution of the parasite, both in the field and in the laboratory (Pigeault et al 2015).

The consequences of *Plasmodium* infections on avian fitness are usually relatively mild but virulence depends on the sensitivity of the host and the parasite lineage. For instance, the accidental introduction of avian malaria into Hawaii played a major role in the decline and extinction of several species of honeycreepers (Atkinson et al. 2000) and still poses a threat to geographically isolated bird species (Lapointe et al. 2012; Levin et al. 2013). Work on wild European bird populations has also revealed strong associations between endemic malaria infection and bird survival and recapture rates (Lachish et al. 2011). More recently, malaria infections have been found to accelerate bird senescence through telomere degradation (Asghar et al. 2015). In addition, some species of avian malaria pose a significant problem to the poultry industry, where mortality rates of up to 90% have been observed in domestic chickens (Springer 1996).

The biology and life cycle of avian malaria parasites in both the vertebrate and vector hosts is similar to that of their mammalian counterparts, but with a few important differences. First, while mammalian parasites have a single exoerythrocytic (EE) cycle in hepatocytes (Huff 1969), avian *Plasmodium* have two obligate exoerythrocytic (EE) cycles, one occurring in the reticuloendothelial system of certain organs, and the other with a much wider tissue distribution (Valkiunas 2004). Second, while certain mammalian parasites (e.g. *P. vivax*) produce dormant forms exclusively during the EE cycle, avian malaria species can also produce dormant forms from the parasite blood stages (Valkiunas 2004). Finally, avian red blood cells are nucleated. Since it could be argued that invasion and growth in nucleated cells-with their richer metabolism and transport-is easier to evolve than development in enucleated mammalian erythrocytes, it is tempting to speculate that these parasites more closely resemble the ancestral state.

In this study we describe the sequence, annotation and comparative genomics of *Plasmodium relictum* SGS1 and *Plasmodium gallinaceum* 8A. Our analyses provide insights into the evolution of unique features of mammalian-infective species and allow an exploration on how far the apparently shared features extend across the entire *Plasmodium* genus. We reveal surprising features involving gene content, gene family expansion and for the first time in *Plasmodium*, the presence of transposable elements.

## Results

### Generation of two avian malaria genomes

Separating parasite and host DNA has been a major obstacle to sequencing avian malaria parasite genomes because avian red blood cells are nucleated. We obtained parasite DNA using two independent strategies (see methods) involving either depleting host DNA based on methylation (Oyola et al. 2013) and using whole genome amplification (*P. gallinaceum*) or sequencing from oocysts from the dissected guts of infected *Culex* mosquitos (*P. relictum*). Using Illumina-sequencing, 23.8-megabase (Mb) and 22.6-Mb high quality drafts of the *P. gallinaceum* and *P. relictum* genomes were produced and assembled into 152 and 498 scaffolds, respectively (Table 1). Both avian *Plasmodium* genomes show very low GC-content; at 17.8 %, *P. gallinaceum* has the lowest GC-content observed in any *Plasmodium* genome sequenced to date.

**Table 1.** Genome statistics for avian malaria genome compared with existing *Plasmodium* reference genomes. Summary data for human infective *P. falciparum* 3D7 (version 3.1) and *P. knowlesi* H (v2, from 01.05.2016) are shown as comparators.

|  | *P. gallinaceum*<br>8A | *P. relictum*<br>SGS1 | *P. falciparum*<br>3D7 | *P. knowlesi*<br>H |
| --- | --- | --- | --- | --- |
| *Nuclear genome* |  |  |  |  |
| Genome size (Mb) | 23.8 | 22.6 | 23.3 | 24.3 |
| N50 | 1,343,538 | 1,833,711 | 1,687,656 | 2,162,603 |
| G+C content (%) | 17.83 | 18.34 | 19.34 | 40.2 |
| Gaps within scaffolds | 1840 | 236 | 0 | 83 |
| No. of scaffolds | 152 | 498 | 14 | 162 |
| No. of chromosomes | ND | 14 | 14 | 14 |
| Amount of N's | 1,156,820 | 27,778 | 0 | 11,431 |
| No. of genes* | 5273 | 5146 | 5429 | 5291 |
| No. of transposable elements (>400bp) | 1244 | 344 | 0 | 0 |
| No. of tRNAs | 46 | 46 | 45 | 45 |
| *Mitochondrial genome* |  |  |  |  |
| Genome size (bp) | 6,747 | 6,092 | 5,967 | 5,957 |
| G+C content (%) | 32.58 | 31.68 | 31.6 | 30.52 |
| No. of genes | 3 | 3 | 3 | 3 |
| *Apicoplast genome* |  |  |  |  |
| Genome size (kb) | 29.4 | 29.4 | 34.3 | 30.6 |
| G+C content (%) | 12.9 | 13.06 | 14.22 | 14.03 |
| No. of genes | 30 | 30 | 30 | 30 |

\* including pseudogenes and partial genes, excluding non-coding RNA genes

The *P. gallinaceum* and *P. relictum* genomes contain 5273 and 5146 genes (Table 1), respectively, (Fig. 1). Those in *P. gallinaceum* were predicted *ab initio* and manually curated using *P. gallinaceum* blood-stage transcriptome data (Lauron et al. 2014) as a guide. These annotated genes were projected onto the *P. relictum* genome, using RATT (Otto et al. 2011) and manually refined.

**Figure 1.**
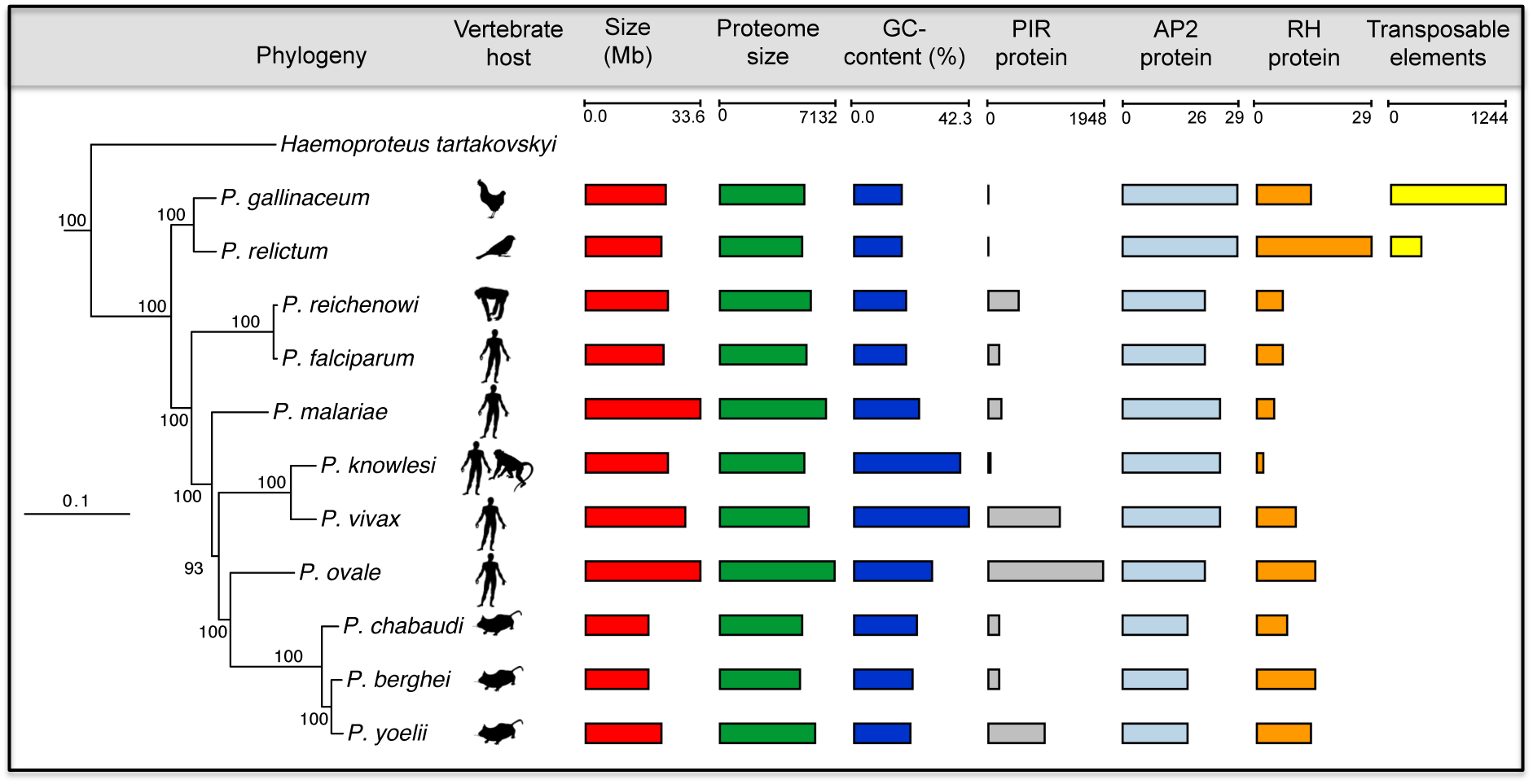
Phylogeny and key features of *Plasmodium* species. Maximum likelihood phylogeny of *Plasmodium* species based on a concatenated alignment of 289,315 amino acid residues from 879 single-copy orthologs. Branch lengths are expected substitutions per amino acid site and values on nodes are number of bootstrap replicates (out of 100) displaying the partition induced by the node. The tree was rooted with sequences from *Toxoplasma* and four piroplasma species, with the full tree shown as Supplementary Fig. S2. The phylogenetic tree is combined with a graphical overview of key features of all reference genomes (genome versions from 1.5.2016). Due to the fragmented nature of the *Haemoproteus tartakovskyi* (Bensch et al. 2016) genome, counts for its key features have not been included.

Excluding subtelomeres, the chromosomes are similar in size and in number of genes to those of other *Plasmodium* species, have positionally conserved centromeres and share synteny across the genus (Supplemental Fig. S1). Likewise, the mitochondrial and apicoplast genomes have been sequenced and show similar size, GC-content and numbers of genes to those previously sequenced from other *Plasmodium* species (Table 1).

### Relationship between *Plasmodium* species

There is broad agreement that three major groups of mammal-infective *Plasmodium* spp are monophyletic but almost every possible arrangement, relative to those that infect birds and reptiles (Blanquart and Gascuel 2011; Perkins 2014) has been proposed at some point. Recently, an extensive multi-locus molecular dataset (Borner et al., 2016) recovered the great ape parasites as the sister group to other *Plasmodium* within a clade of mammalian parasites, disagreeing with the earlier phylogenomic analyses (Pick et al. 2011) that had supported the hypothesis (Waters et al. 1991) that mammalian *Plasmodium* are polyphyletic, and that *P. falciparum* and its relatives evolved recently from an avian ancestor, perhaps explaining its high virulence.

We re-evaluated the phylogeny of the mammalian groups with genome-wide data using both Bayesian and maximum-likelihood models. We found robust support for *P. gallinaceum* and *P. relictum* forming an outgroup to the other *Plasmodium* species, and the *Laverania* appearing as the sister group to other mammalian *Plasmodium* (Fig. 1; Supplemental Fig. S2). We also found that subgenus *Plasmodium* is paraphyletic, with an unexpected sister-group relationship between *P. ovale* and the rodent-infective species and *P. malariae* (Rutledge et al. 2017) branching as the deepest lineage outside the avian species and the *Laverania*. This result is robust to changes in the substitution model used for phylogenetic inference (see Supplemental information, Supplemental Fig. S3A).

Correctly placing the wider outgroup of more distantly related *Apicomplexa* is a widely recognized difficulty (Perkins 2014; Borner et al. 2016). Our attempts to fit more complex and potentially more realistic phylogenetic models to resolve discrepancies between Bayesian and maximum likelihood trees were unsuccessful, as MCMC runs failed to converge. However, our data did give strong and consistent support for the relationships within *Plasmodium* and for the root of *Plasmodium* when the data for *Haemoproteus* were used as an outgroup (Supplemental Fig. S3B, C). Sparse molecular data are available for many lineages of the genus *Plasmodium* (Perkins and Schaer 2016) and molecular data from these and additional *non-Plasmodium* lineages of *Haemosporidia*, especially those closely related to mammal and bird *Plasmodium*, will be key to fully resolving the evolution of the human pathogens.

### Dating speciation across the *Plasmodium* genus

Dating speciation events in the *Plasmodium* lineage has been controversial and hindered by a lack of fossil records. However, the availability of multiple genomes for several lineages has enabled coalescence based methods to provide revised timings (Rutledge et al. 2017). Using dates from the latter, we calibrated amino acid divergence across the tree to approximate speciation times (Fig. 2), as previously described (Silva et al. 2015), with the caveat that the dates assume equal rates of divergence across all branches. *P. gallinaceum* and *P. relictum* appear to have diverged 4 million years ago and that the avian lineage arose, along with the radiation of all sequenced *Plasmodium* species, around 10 million years ago, much more recently than the avian-mammalian split around 300 million years ago (Kumar and Hedges 1998). This result is consistent with recent data from the *Laverania* subgenus that shows parasite speciation events are much more recent than that of their hosts (Otto et al. 2017).

**Figure 2.**
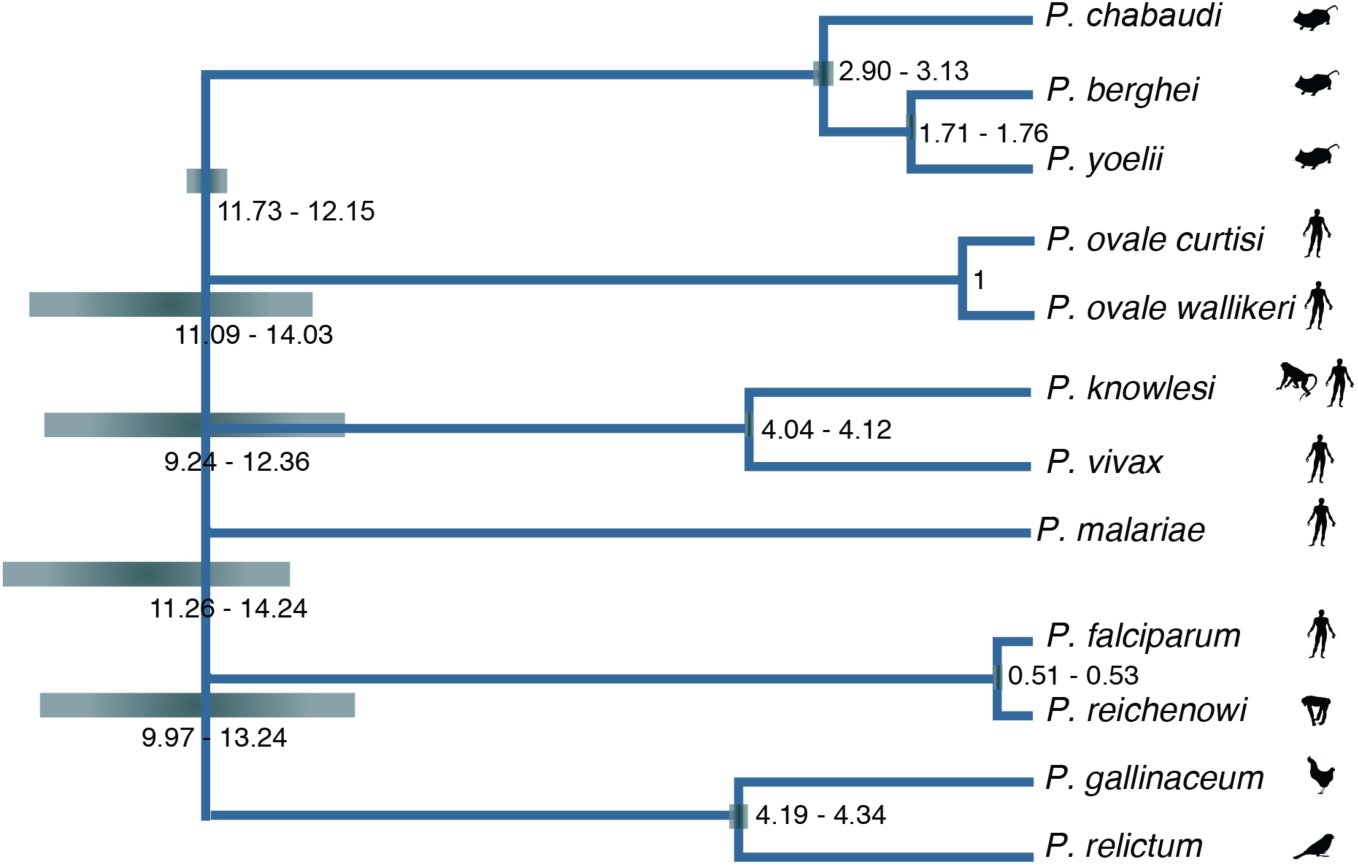
Schematic of the phylogenetic tree showing approximate speciation times across the *Plasmodium* genus. Species dates were estimated using a total Least Squares regression on the dAA values (Silva et al. 2015) and calibrated on the split of two *P. ovale* species, which is assumed to have occurred one million years ago (Rutledge et al. 2017). 95% confidence intervals for each node are represented by heat maps.

### Novel genomic features of the avian lineage

We found 50 avian malaria-specific core genes with orthologs in both species (Supplemental Table S1). Using the available *P. gallinaceum* transcriptome data (Lauron et al. 2014), we found that only two of these 50 genes showed evidence of expression in the blood stage indicating that the remaining 48 genes are likely to play a role elsewhere in the life cycle. For the majority (52%) of these genes, putative functions could not be ascribed (Supplemental Table S1) but a possible new member of the AP2 family of transcription factors (PRELSG 1134000, PGAL8A 00142800) (Supplemental Fig. S4A) was found in both species. We also found a 6-cysteine protein, a protein phosphatase and an AMP-specific ABC transporter. To date, analyses have failed to identify homologs in *Plasmodium* of the non-homologous end joining (NHEJ) pathway that repairs double-strand breaks in DNA. Ku70 is a member of this pathway that has apparent orthologs in both *P. gallinaceum* and *P. relictum* (PGAL8A_00014200, PRELSG_0411800), supported by a three-dimensional model created using I-TASSER (Yang et al. 2015) (Supplemental Fig. S5). However, an ortholog of Ku80 (the obligate partner of Ku70 in NHEJ activity) is not present (Fell and Schild-Poulter 2015).

An unusual SNF1-like kinase, KIN, plays a key role in cellular energy metabolism in *Plasmodium* spp. (Mancio-Silva et al. 2017). KIN functions as a single subunit, unlike the trimeric structure of canonical AMP-activated kinases. Unique to the avian malaria genomes is evidence of a more typical SNF1-like kinase; in *P. gallinaceum* and *P. relictum*, alpha (PGAL8A_00159300, PRELSG_1117500) and beta subunits (PGAL8A_00165250, PRELSG_1111850) can be clearly identified based on domain analysis. Based on an analysis of Pfam domains and structural prediction using I-TASSER, we were also able to find a possible candidate gamma subunit in each of the two avian malaria species (PGAL8A_00033950, PRELSG_1019550).

The core regions of *P. relictum* and *P. gallinaceum* chromosomes have the same complement of genes except for a putative gene of unknown function in *P. relictum* (PRELSG_0909800) (Supplemental Table S1, Supplemental Fig. S4B) and several differentially distributed pseudogenes (Supplemental Table S1, Supplemental Fig. S6).

We found 15 genes present in *P. gallinaceum* and *P. relictum* that were previously defined as *Laverania* specific (Supplemental Table S2). Apart from hypothetical proteins this includes ATPase1 (Supplemental Fig. S7A), apyrase and a sugar transporter. We also found 12 genes that have not previously been identified outside the *vivax, ovale or malariae* clades (Supplemental Table S3). Among these are the merozoite surface protein 1 paralog (MSP1P) and an ApiAP2 transcription factor (Supplemental Fig. S7B).

The shikimate pathway provides precursors for folate biosynthesis but is remarkably different between mammalian and avian *Plasmodium* spp. In the latter, genes encoding two enzymes in the pathway appear to have become pseudogenes (Supplemental Fig. S8, Supplemental Table S4) and the gene encoding a key enzyme complex, the pentafunctional AROM polypeptide, is completely missing. Thus avian malaria parasites are not able to synthesize folate *de novo*. One explanation for this could be the fact that the host cells are nucleated and therefore provide a richer nutrient environment. Three other core genes are missing in *P. gallinaeum* and *P. relictum* (Supplemental Table S5) in addition to AROM, but all three are conserved hypothetical proteins.

### A family of Long Terminal Repeat (LTR)- Retrotransposons in avian malaria genomes

Despite the presence of retrotransposons in the majority of eukaryotic genomes, none have yet been identified from *Plasmodium* species. Surprisingly, we identified a large number of transposable element (TE) fragments in the avian malaria genomes (Supplemental Fig. S9): 1244 in *P. gallinaceum* and 344 in *P. relictum* (Table 1). The vast majority (Supplemental Fig. S10, Supplemental Fig S11A) were found in the subtelomeres (Supplemental Fig. S9, Fig. 3C). A single complete, 5.7 kb retrotransposon is present in *P. gallinaceum* (PGAL8A_00410600) (Fig. 3A) and contains a 4.5 kb open reading frame encoding a gag-pol polyprotein including the following domains: a retroviral aspartyl protease (Pfam:PF00077), reverse transcriptase (Pfam:PF00078), RNase H (Interpro:IPR012337) and integrase (Pfam:PF00665). It is bounded by long terminal repeats of 459 nucleotides (5’LTR) and 469 nucleotides (3’LTR) respectively and contains a primer binding site and polypurine tract (Fig. 3B). Based on the order of encoded HMM domains, the *P. gallinaceum* retrotransposon can be classified as Ty3/Gypsy retrotransposon (Steinbiss et al. 2009). In addition to the complete TE, we found four nearly full-length copies, also bounded by long terminal repeats (PGAL8A_00328600, PGAL8A_00325400, PGAL8A_00189500, PGAL8A_00270200) (Fig. 3A). *P. relictum* did not contain a complete retrotransposon but based on the programs LTRharvest/LTRdigest (Ellinghaus et al. 2008; Steinbiss et al. 2009), we found 7 near full-length copies with all the required domains. The most complete is localized in the core area on chromosome 6 (Supplemental Fig S11B). It has a length of 5.3 kb, contains all the required HMM domains and is bounded by long terminal repeats of 253bp (5’LTR) and 257 bp (3’LTR) that are shorter than those observed in *P. gallinaceum*. A BLAST comparison showed highest similarity (28%) to a retrotransposon described in *Ascogregarina taiwanensis*, a gregarine that infects mosquito larvae (Templeton et al. 2010). This is also reflected in the phylogenetic tree (Supplemental Fig. S12).

**Figure 3.**
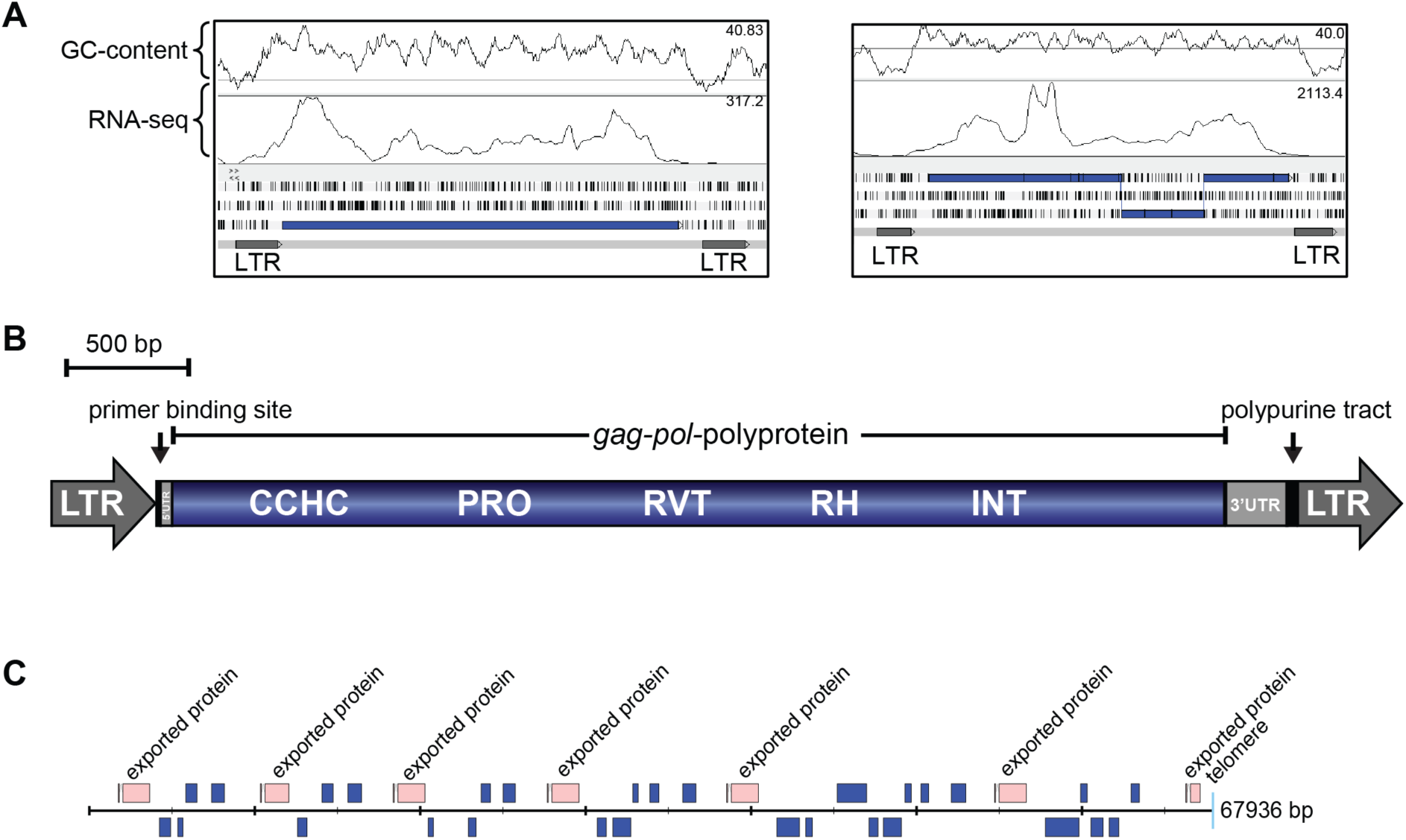
Transposable elements in *P. gallinaceum*. (A) Artemis screenshot showing a complete retrotransposon of *P. gallinaceum* (PGAL8A_00410600) and a copy where the open reading frame encoding gag-pol-polyprotein is frame-shifted. (B) Diagram of the *P. gallinaceum* retrotransposon (PGAL8A_00410600). The Ty3/Gypsy transposable element contains a continuous open reading frame including a CCHC-type zinc finger domain (CCHC), aspartic protease domain (PRO), reverse transcriptase domain (RVT), RNase H domain (RH) and an integrase domain (INT). The element is bounded by long terminal repeats (LTR). (C) A single subtelomeric region (contig 70) from *P. gallinaceum*. Transposable elements are shown in blue.

Despite their high degree of fragmentation, we were able to align 71 regions of ≥2kb common to the TEs of both species, resulting in a trimmed tree of 69 sequences aligned over 1295bp. The resulting phylogenetic tree suggests at least 2 independent acquisitions in the two different species (Fig. 4). Within *P. gallinaceum* there are two major clades and that can also be differentiated based on GC content. Older sequences would be expected to converge to the level of the endogenous genome, which in the case of the avian malaria is extremely low (Table 1). There is evidence of TE transcription in existing RNA-seq data (Fig 3A) but it is not possible to map the data meaningfully to individual TE copies due to unevenness of coverage and a paucity of discriminating SNPs.

**Figure 4.**
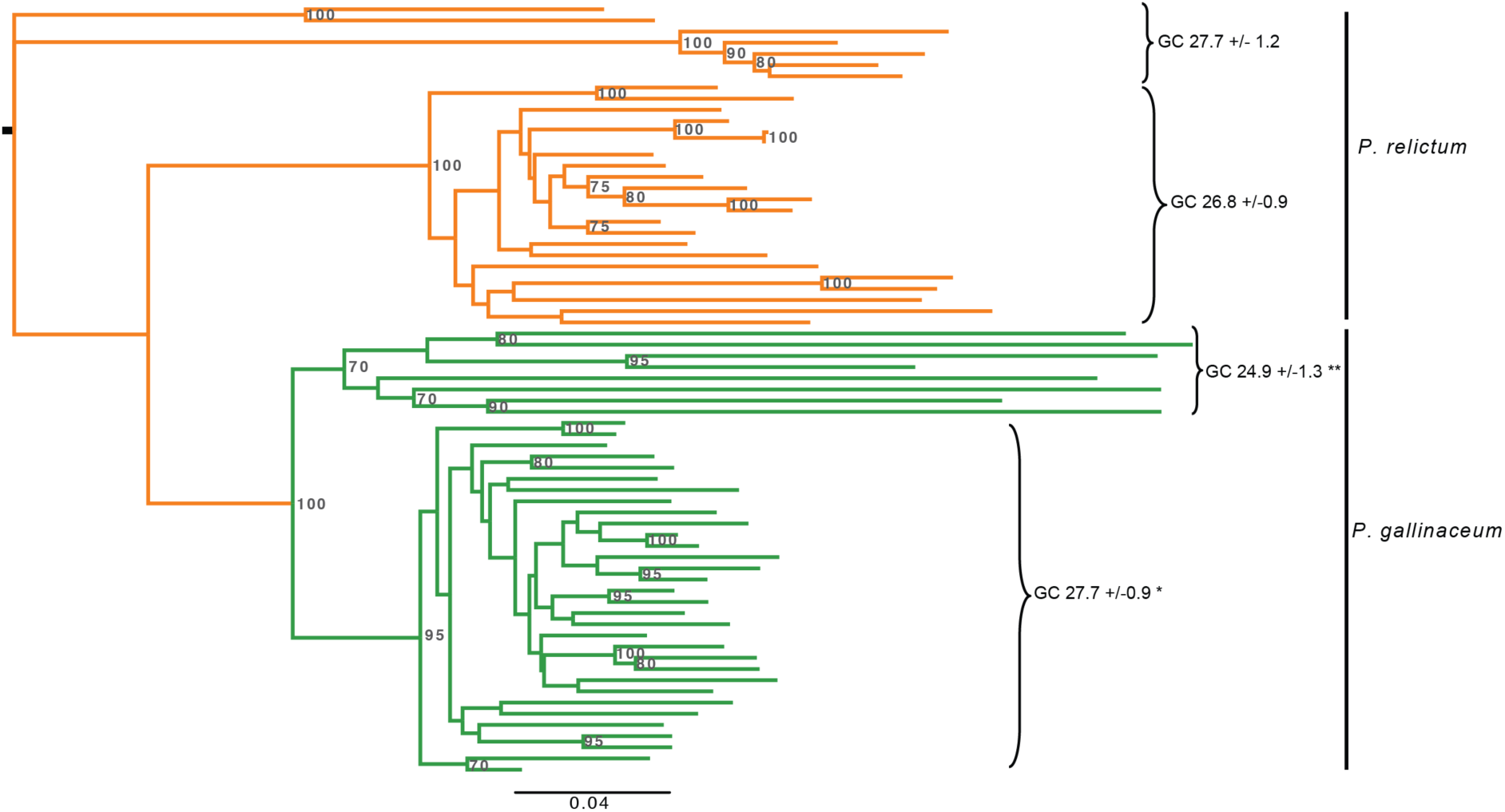
Phylogenetic analysis of 69 transposable elements from *P. gallinaceum* and *P. relictum*. For each element, GC-content is shown and clearly distinguishes two clades in *P. gallinaceum*. Unrooted maximum likelihood tree based on nucleotides using the GTR+G evolutionary model. Bootstrap values <70 are not shown. Percentage GC values indicate mean +/- variance. P-values were determined based on a simple randomisation approach (*P=<0.01, ** P=<0.0001) see Supplemental Methods.

To test the activity of the complete retrotransposon, we attempted to introduce a gag-pol expression cassette into the rodent malaria parasite *P. berghei*. Transfection was attempted on four independent occasions with no integration of the *P. gallinaceum* gag-pol expression cassette being detected. Parallel transfection of a second vector containing an unrelated insert acted as a positive control, ruling out technical difficulties. Failure to introduce the *P. gallinaceum* gag-pol transposase expression cassette is interpreted as potential toxicity associated with expression of the *P. gallinaceum* gag-pol under the very strong *pbhsp70* promoter and attempts to swap the promoter for the weaker *pbeef1a* promoter, or an inducible promoter are ongoing.

### Multigene families

In addition to the multigene families present in previously sequenced *Plasmodium* genomes e.g. ETRAMPS, *pir* and reticulocyte binding proteins (Gardner et al. 2002; Pain et al. 2008), (Table 2, Supplemental Table S6), we identified four novel gene families in the avian malaria genomes (Supplemental Fig. S13) and found that a *Plasmodium* specific low copy number gene is expanded in the avian species. To maintain consistency with the gene family naming scheme established for other species (Otto et al., 2014a), the families are named *fam-e* to *fam-i* (Supplemental Fig. S13).

**Table 2.** Members of subtelomeric multigene families in the genomes of *P. gallinaceum* 8A, *P. relictum* SGS1

| Gene family | No. of gene members |  |
| --- | --- | --- |
|  | *P. gallinaceum* | *P. relictum* |
| **PIR-like protein** | **20** | **4** |
| PIR-like, pseudogene | 1 | 0 |
| PIR-like, fragment | 1 | 1 |
| **surface-associated interspersed protein (SURFIN)** | **40** | **14** |
| surface-associated interspersed protein (SURFIN), pseudogene | 4 | 16 |
| surface-associated interspersed protein (SURFIN), fragment | 35 | 20 |
| **early transcribed membrane protein** | **12** | **12** |
| early transcribed membrane protein, pseudogene | 0 | 1 |
| early transcribed membrane protein, fragment | 0 | 1 |
| **reticulocyte binding protein, putative** | **8** | **13** |
| reticulocyte binding protein, pseudogene | 4 | 5 |
| reticulocyte binding protein, fragment | 2 | 15 |
| **fam-e protein** | **38** | **4** |
| fam-e protein, pseudogene | 0 | 0 |
| fam-e protein, fragment | 11 | 0 |
| **fam-f protein** | **16** | **14** |
| fam-f protein, pseudogene | 2 | 0 |
| fam-f protein, fragment | 0 | 1 |
| **fam-g protein** | **107** | **0** |
| fam-g protein, pseudogene | 2 | 0 |
| fam-g protein, fragment | 12 | 0 |
| **fam-h protein** | **2** | **49** |
| fam-h protein, pseudogene | 0 | 0 |
| fam-h protein, fragment | 0 | 0 |
| **fam-i protein** | **23** | **0** |
| fam-i, pseudogene | 0 | 0 |
| fam-i, fragment | 3 | 0 |

To explore the relationship between *Plasmodium* subtelomeric gene families across the genus we used two different clustering approaches, based either on global similarity or on conservation of short motifs. First, we compared all genes with BLASTP and created a gene network, where the genes (nodes) were connected if they shared a global similarity above a threshold (Fig. 5A). Although the topology of the network changed with different sequence identity thresholds (Supplemental Fig S14A), at a threshold of 31% the STP1 and the Surface-associated interspersed proteins (SURFINs) of different species are connected. The SURFINs are encoded by a family of 10 genes in both *P. falciparum* and *P. reichenowi*. We found a relatively high number of SURFINs in avian malaria genomes, 40 in *P. gallinaceum* and 14 in *P. relictum* (Table 2). As shown in previous studies, SURFINs show some sequence similarity to PIR proteins of *P. vivax* (Merino et al. 2006; Winter et al. 2005) and some SURFINs share a domain with the SICAvars of *P. knowlesi*. To examine this relationship more closely by highlighting similarity that could be missed by BLASTP, we used MEME (Bailey et al. 2009) to generate 96 sequence motifs from the STP1 and SURFIN families, respectively. Next we searched for those predicted motifs in all predicted proteins (excluding low complexity regions) of the 11 sequenced *Plasmodiu* species and visualized the results as a binary occurrence matrix (Supplemental Fig. S14B). Although some proteins share a limited repertoire of the STP1 or SURFIN motifs-namely DBL containing protein, antigen 332 and three putative proteins of unknown function (PRELSG_1445700, PmUG01_00032900, PmUG01_10034200), we observed extensive motif-sharing amongst the STP1 and SURFIN proteins (Fig. 5b) but only a single motif is shared between STP1, SURFIN and SICAvar (Supplemental Fig. S14B,C). This suggests that STP1 and SURFIN comprise a superfamily. The SURFINs cluster into two groups. Group II is unique to *P. gallinaceum* but group I includes both homologs from the avian *Plasmodium* and the hominoid *Laverania* subgenus. The STP1 proteins are not found in the avian malaria parasites but form two *P. ovale* and one *P. malariae* specific clusters. Whether these poorly characterized families have functional similarities remains to be determined.

**Figure 5.**
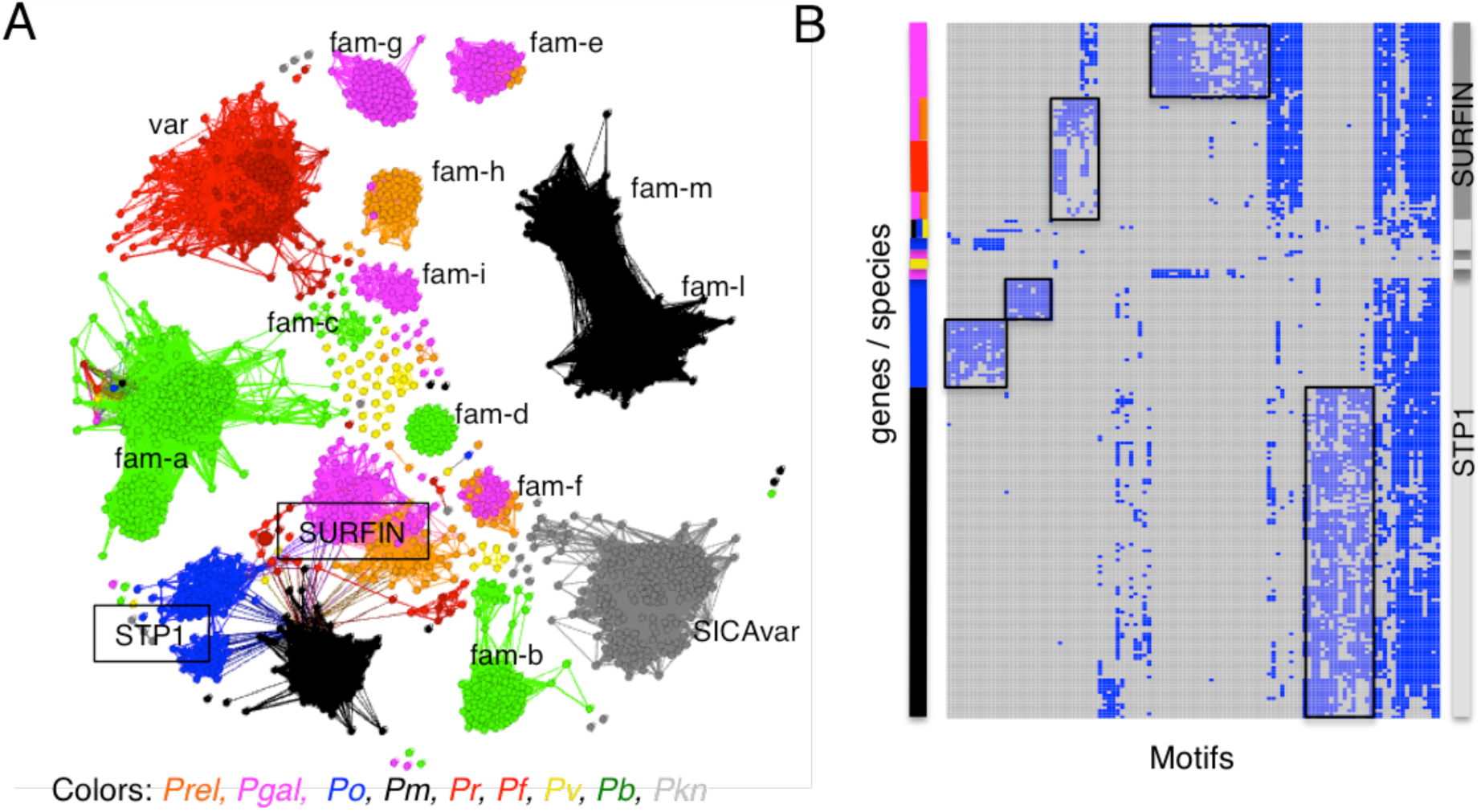
Similarity of gene families within *Plasmodium*. (A) A network of BLASTp similarity between genes (nodes) sharing at least 31% global identity. Genes are coloured by species. The *pir* genes were excluded due to their large numbers across the *Plasmodium* genus. *Fam-m* and *Fam-l* are *P. malariae* specific gene families (Rutledge et al. 2017) (B) Clustering of STP1 and SURFIN genes based on the occurrence motifs identified using MEME. Where a gene (row) has a specific motif (column) the value is set to one. The matrix is clustered through a hierarchical clustering algorithm (Ward 1963), to visualize similar patterns of motif-sharing. The x-axis represents motifs that occur in at least 10 genes, and individual genes are displayed on the y-axis (rows). Coloured bars on the left identify species, the bar on the right the gene annotation. Boxed areas indicate possible gene family sub-types.

The *pir (Plasmodium* interspersed repeat) genes are the largest multigene family in *Plasmodium* spp. and have been found in high numbers in all malaria species sequenced to date (Janssen et al. 2004). In the avian malaria genomes, we found only a small number of distantly related genes that are possibly members of the family: 20 in *P. gallinaceum* and 4 in *P. relictum* (Table 2). They follow the canonical 3-exon structure, with the second exon encoding a cysteine-rich low-complexity sequence, a transmembrane domain and a highly conserved third exon. However, the avian *pir* genes have only remote sequence similarity to those of other *Plasmodium* species (and have therefore been annotated as *pir*-like); the highest sequence similarity (41% over 60 amino acids) was found between a *pir* from *P. vivax* (PVP01_0800600) and *P. gallinaceum*.

We identified 290 genes in *P. gallinaceum*, and 203 in *P. relictum*, encoding the PEXEL motif that is frequently present in *Plasmodium* sub-telomeric multi-gene families and is important for trafficking proteins into and through the host cell (Marti et al. 2004). Three families in particular appear to be important in the avian malaria lineage. The *fam-f* family has only a single member in each species of the *Laverania* subgenus (PF3D7_1352900, PRCDC_1351900) and two members in the *vivax* clade (PVP01_1201900, PVP01_1147000, PKNH_1248100, PKNH_1148800) but has 16 members in *P. gallinaceum* and 14 members in *P. relictum* (Supplemental Fig. S13, Supplemental Table S6). Using I-TASSER to predict the structure, *fam-f* has some similarity (a modest C-score of −2.33) with human alpha catenin (PDB:4IGG) that is involved in cell adhesion. To date, *fam-g* and *fam-h* appear to be specific to avian malaria. *Fam-g* has 107 members in *P. gallinaceum* and none in *P. relictum*, whereas *fam-h* has 49 copies in *P. relictum* and only 2 in *P. gallinaceum* (Supplemental Table S6). It is possible that the relative absence of *pir* genes might be in some way compensated by the expansion of these families.

There are two additional novel gene families in the avian malaria genomes, *fam-e* and *fam-i. Fam-e* is present in both avian malaria species and is a 2-exon gene with an average length of 350 amino acids and a transmembrane domain. There are 38 copies in *P. gallinaceum* and only 4 in *P. relictum* (Table 2, Supplemental Table S6). *Fam-i* is a 2-exon gene family only present in *P. gallinaceum* with 23 members (Supplemental Fig. S13). Both gene families lack a putative PEXEL motif. One aspect of these gene families is that the expression of a few individual members seems to dominate within the asexual blood stages, see Supplemental Table S7.

### Expansion of the reticulocyte-binding protein (RBP) family in avian malaria parasites

Homologs of reticulocyte-binding protein (RBP) are important in red cell invasion yet a recent publication (Lauron et al. 2015) indicated, based on a transcriptome assembly, a lack of RBPs in *P. gallinaceum*. In contrast, we found an expansion of this family in both avian malaria parasites with 8 copies in *P. gallinaceum* and at least 29 in *P. relictum* (33 if fragments are included; see Methods). Because these genes are long (> 7.5 kb) and have large blocks of high sequence similarity they are difficult to assemble in *P. relictum* and the copy number in this species could be underestimated. However, we can see in a maximum likelihood tree of avian RBP ≥ 4.5kb that the separation into different sub-families predates the speciation of the two avian malaria lineages (Supplemental Fig. S15A). As with the STP1 and SURFIN families, we analysed sequence motifs produced by MEME to investigate the relationship and evolution of the RBPs in nine *Plasmodium* spp (Fig. 6A, Supplemental Fig. S15B, C) in more detail. Conserved sequences were used, corresponding to two sets of shared motifs (black dashed box and stars in Fig. 6A), to draw maximum likelihood trees for the nine *Plasmodium* spp (Fig. 6B). The phylogenetic analysis shows that the genes predate the speciation of *Plasmodium* genus but we see a strong host-specific diversification. The RBPs of *P. ovale, P. malariae, P. vivax* and *P. knowlesi* form three different clades. Interestingly the *P. berghei* RBPs seem to be very similar to each other, but very different to the other species. This general classification of the species can also be seen in the motif occurrence matrix (Fig. 6C). Some of the motifs are shared in all RBP. We also find host specific motifs, splitting the rodent, avian and human and primate hosts.

**Figure 6.**
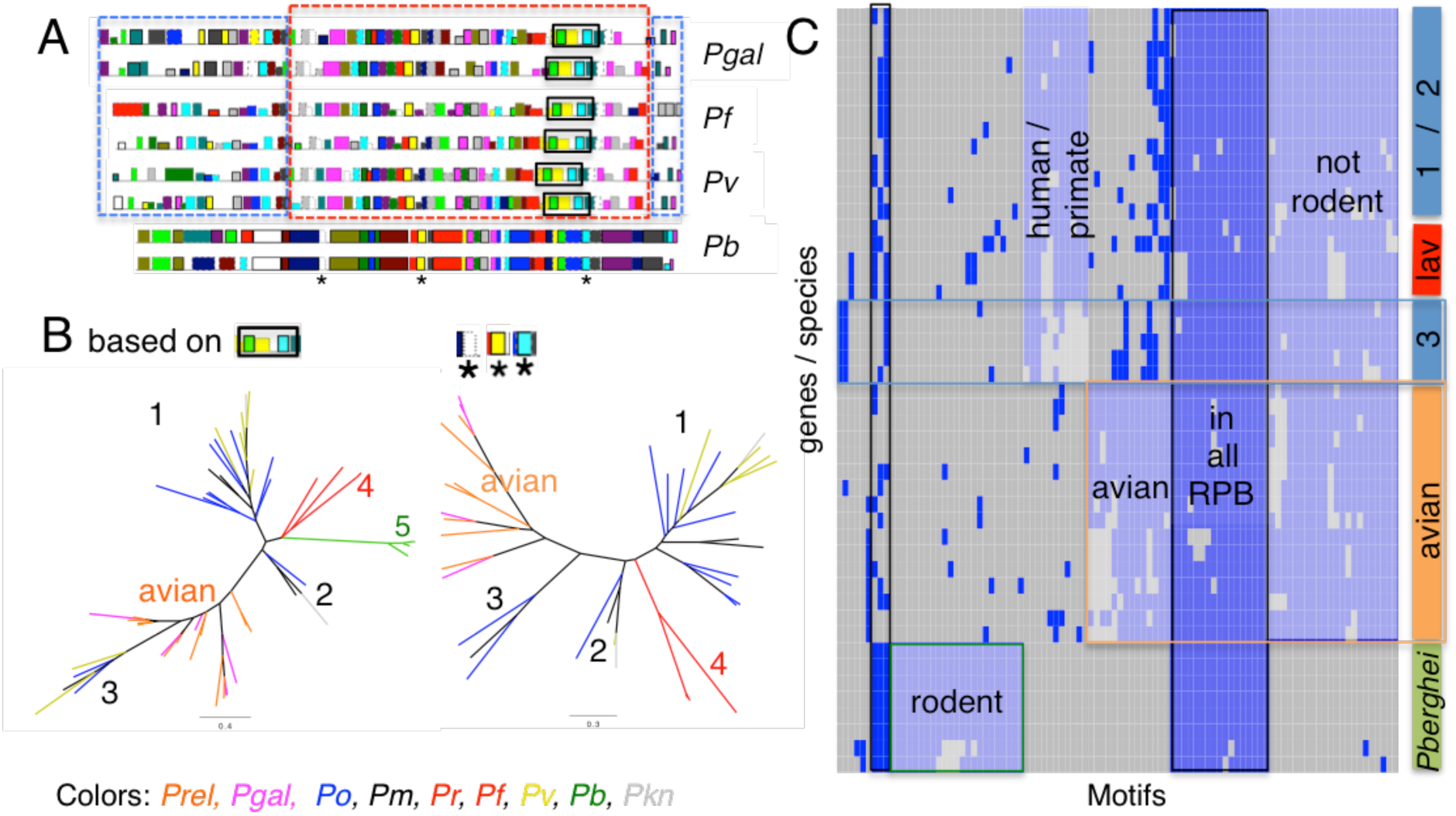
RBP Meme motifs comparison. Analysis of 96 meme motifs obtained from reticulocyte binding proteins (RBP) of nine species. (A) Example of motifs predicted on two RBP from each of four species. Each colored rectangle (along the protein) represents a different one of the 96 motifs, with their heights corresponding to their respective e-values. The red dashed box around the sequences of *P. gallinaceum, P. falciparum* and *P. vivax* highlights a similar order of motifs. The blue dashed boxes on either side highlight differences in motif content. The black box and the three stars are motifs used to build the tree in (B). (B): Two maximum likelihood phylogenetic trees based on two motif sets. The left tree was generated using the three motifs (indicated with an asterisk * in Panel A, in total 72 amino acid long) and the second tree was generated using the motifs from the black box in panel A, 169 amino acids long (all bootstrap values are 100). Labels 1, *2* and 3 identify the distinct clusters of the *P. malariae, P. ovale* and *P. vivax* RBP, as previously reported (Rutledge et al. 2017), 4 *P. falciparum* and *P. reichenowi* and 5 *P. berghei*. (C) Clustering of the binary occurrence of meme motifs for each RBP, similar to Fig. 5B. The bar on the right represents either species (lav-*Laverania*, avian, *P.berghei*) or the groups 1,2 and 3 from (B) This analysis does not split group one and two of *P. malariae, P. ovale* and *P. vivax* RBPs. The x-axis represents the 96 motifs. Blue represents at least one occurrence of that motif for that gene. Shared patterns are highlighted with coloured boxes.

### Lineage-specific diversification

Using pairwise *d_N_/d*_S_comparisons in PAML (Yang 2007), we looked for signatures of selection across 4285 orthologous genes. Between the major clades, *d*_S_clearly saturates (indicated by extremely low *d*_N_ */d*_S_values; Supplemental Table S8). We therefore focused on within clade comparisons -*P. gallinaceum* with *P. relictum, P. falciparum* with *P. reichenowi* and *P. vivax* with *P. knowlesi* - and took the top 250 *d_N_/d*_S_values from each pairwise comparison. Across all comparisons, the list of 188 annotated genes (Supplemental Table S9) contains those with known important links with parasite biology (host invasion). In addition, there is an enrichment for genes of unknown function (68% compared with <40% in the whole genome) suggesting unexplored areas of parasite biology. Just 28 genes were common to the top 250 of all three within-clade comparisons and 22 of these were also of unknown function. The remaining six, again reflect highly characterized functions that are important for parasite biology (Supplemental Table S9). Across all of the comparisons, the only significant enrichment occurred in the comparison of avian *Plasmodium* genes and involved the term ‘entry into host’.

To assess diversification across the genus, in the face of the saturated *d_S_* values, we identified the top 250 *d_N_* values in comparisons within and between the three major lineages (Supplemental Table S10), containing *P. gallinaceum, P. falciparum* and *P. vivax*, respectively. Surprisingly across the entire analysis, there is a clear enrichment for uncharacterized genes (P<0.0001, two tailed Fisher’s exact test). The only genes to show significantly enriched functional annotation were those that were either unique to both the avian and falciparum clades or the avian clade alone (Fig. 7), and were restricted to ‘symbiont-containing vacuole membrane’, ‘rhoptry’ and ‘entry into host cell’, confirming the striking adaptation of host entry genes in the avian lineage.

**Figure 7.**
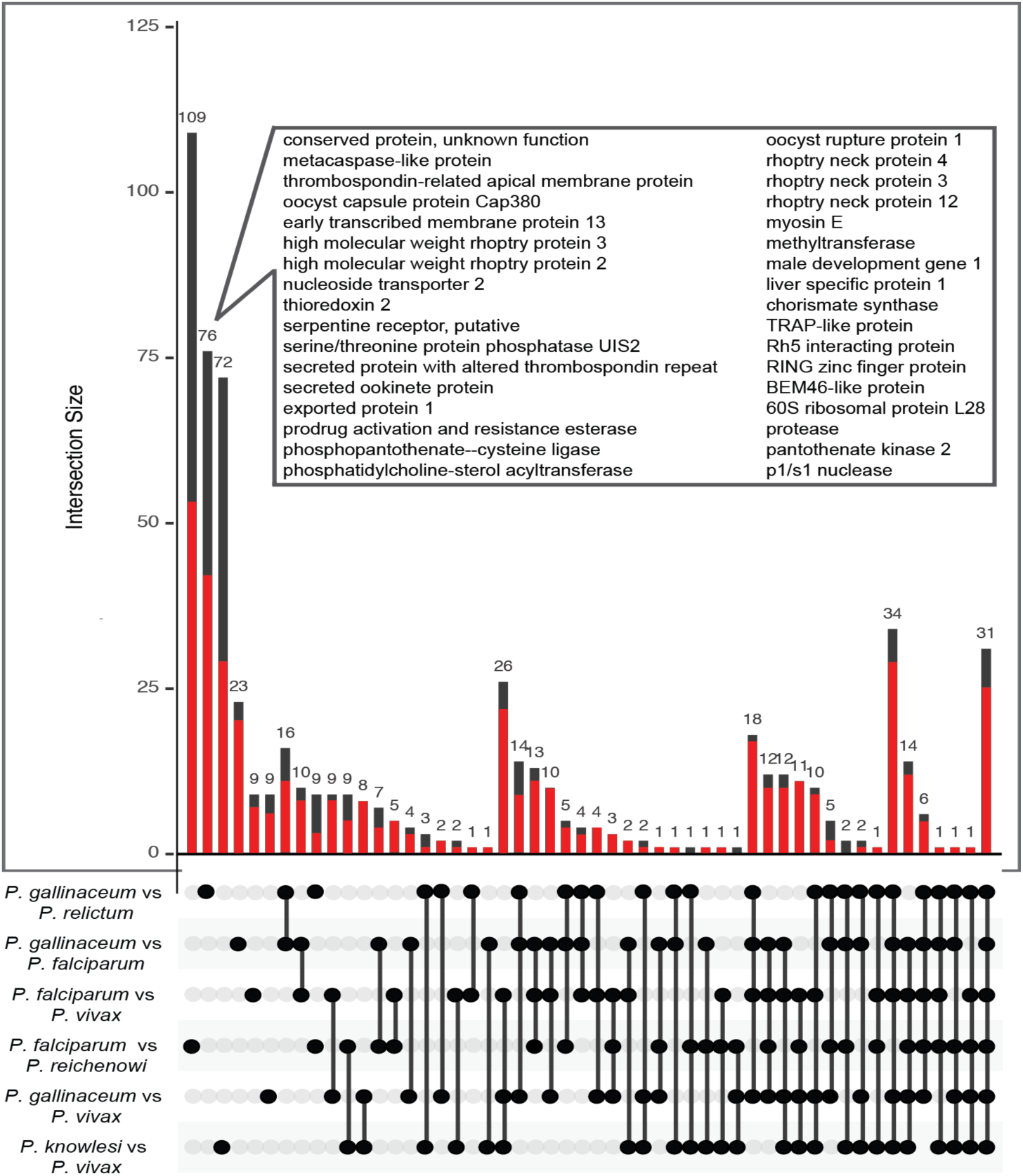
Analysis of genes with high rates of non-synonymous substitutions (*d*_N_) between six species. From pairwise comparisons within- and between-clades, the 250 highest scoring genes were selected. The matrix shows the intersections between the six gene lists and the barplot above shows the number of genes that are unique to each intersection. The fraction of genes with unknown function in each category is shown with a red bar. The gene products are shown for the avian species comparison, which had the most significant GO term enrichment.

## Discussion

Until now, all high quality and manually curated malaria genomes have been from mammalian parasites. The reference genome assemblies of two avian malaria genomes, *P. gallinaceum* and *P. relictum*, in the present study occupy a sister group relative to the phylogeny of the mammal-infecting *Plasmodium* species. To confirm this sister group was challenging, as the outgroups are diverse in sequence and the avian parasites share the same extreme GC bias (19%) as those from the *Laverania* subgenus. All other mammalian *Plasmodium* genomes have a GC content, between 23-44% and it has been suggested that this difference is due to the lack of efficient base excision repair (BER) (Haltiwanger et al. 2000) that drives the genome towards lower %GC content. If this is the case, it is likely that BER has been lost in both the *Laverania* and avian malaria lineages or that improvements to BER occurred after the evolution of the *Laverania* branch.

We have analysed 11 parasite species from diverse hosts and nearly all genes in regions previously defined as a conserved core occurred as 1:1 orthologs. The roles of most genes are therefore probably shared between the species and transcend host differences. Our analysis therefore focused on genes that are not shared to investigate species-specific malaria biology. For example, we find 50 core genes (1:1 orthologs) that are unique to the avian malaria parasites. As these include a novel AP2 gene, encoding a class of transcription factor known for its importance in developmental regulation, and the majority (48) of the unique genes are not expressed in blood stages, it is possible that they play a role in the second EE cycle unique to avian species (Garnham 1966). The differences in the folate and heme pathways of the avian species could also be attributed to their colonization of the more metabolically competent nucleated red blood cells of birds.

In the sub-telomeres we discovered four new gene families (*fam-e, fam-g, fam-h* and *fam-i*), a newly expanded family *fam-f*, expansion of SURFIN types and a reduction in the number of *Plasmodium* interspersed repeat (*pir*) genes compared to other species. *pir* genes are present in all *Plasmodium* species sequenced to date and are the largest multi-gene family within the genus. Their function is unclear, but their protein products are present at the host parasite interface, are implicated in parasite-host interactions (Goel et al. 2015; Niang et al. 2014) and have been associated with immune evasion (Cunningham et al. 2010; Fernandez-Becerra et al. 2009; Saito et al. 2017). In the *P. chabaudi* rodent model, differential *pir* gene expression is associated with parasite virulence (Spence et al. 2013). How avian parasites replace this function is unclear. *fam-f* is single copy in the mammalian malaria genomes but is significantly expanded in both avian genomes and shows a distant homology to a human protein that is involved in cell-adhesion. SURFINs are also found on the surface of infected red blood cells, are expanded in *P. gallinaceum* and show substantial similarity to the STP1 family which leads us to hypothesise that these two genes families may have shared a common ancestor and their sequences evolved in a host-dependent manner.

Another polymorphic gene family involved in host cell invasion, and recently attributed to host specificity (Otto et al., 2014b) and red blood cell preference, are reticulocyte binding proteins (RBPs). This family is significantly expanded in *P. relictum*, which could explain the ability of this parasite to infect a wide range of avian species and tissues. We also see strong host-specific diversification that in combination with the function of “entry into host” being enriched in several comparisons suggests that the optimization of these genes to their host environment is of evolutionary importance. The rodent RBP cluster together and are different to the other clades but still share certain motifs across the genus (Fig. 5A). Interestingly, the more diverse motifs are found at the N-terminus of the RBP, Fig. 6A, (blue dashed boxes). The differences between the N-termini of the RBP, is intriguing as these regions mediate binding to host receptors. It is tempting to speculate that the conserved motifs are important for the general structure of the RBP, but the more variable N-terminal regions evolved to bind to specific host receptors.

The most striking difference between the avian parasites and their mammalian infecting relatives is the presence of long terminal repeat (LTR) retrotransposons. The only other retrotransposon found so far in *Apicomplexa* are those from *Eimeria* (Ling et al. 2007; Reid et al. 2014). Both the retrotransposons found in *Eimeria* and the ones in the avian malaria parasites belong to the Ty3/Gypsy family. The transposable element found in *Eimeria* is similar to chromoviruses, a subgroup of Ty3/Gypsy retrotransposons, whereas the TE from *P. gallinaceum* does not contain chromodomains. Both TEs therefore seem to be from different lineages. We were able to identify several unreported fragments of Ty3/Gypsy retrotransposons in the recently published genome of the bird parasite *Haemoproteus tartakovskyi* (Bensch et al. 2016), a sister genus of *Plasmodium*. Although the TE was found in three avian parasite species, it appears that these were independent acquisitions. Moreover, in *P. gallinaceum* there appear to be two distinct radiations of TEs that can be differentiated based on their branch lengths and GC content. The two radiations may therefore represent two temporally distinct introductions that differentially equilibrated to the GC content of the host genome overtime. The similarity of the avian *Plasmodium* TE to a sequence from the vector parasite *Ascogregarina taiwanensis* suggests that the TEs within the avian *Plasmodium* species were horizontally acquired from vectors that may have been co-infected with *Ascogregarines*. However, the question remains why transposable elements (TEs) were not found in any other *Plasmodium* species sequenced to date. With multiple acquisitions into the avian-infective lineage but none in any other lineage, avian *Plasmodium* must therefore be either more exposed or more permissive to TEs.

To date, *Piggybac* is the only TE to have been successfully mobilised in *Plasmodium* under experimental conditions (Balu et al. 2009). We have made multiple attempts to express *P. galllinaceum* gag-pol in *P. berghei* but these have been unsuccessful, perhaps due to its toxicity. Understanding the mechanism of action for this novel TE could open up exciting new possibilities for TE based insertional mutagenesis within *Plasmodium* species.

Availability of complete genome data allowed parasite evolution to be examined across the mammalian and avian clades of *Plasmodium*. Signatures of diversifying selection in host-interacting genes have previously been uncovered in the *P. falciparum* and *P. vivax* lineages (Otto et al. 2014; Neafsey et al. 2012) However, what was surprising, was the extent to which invasion genes have diversified in the avian lineage, possibly reflecting the increased complexity in the avian parasite life cycles, with two extraerythrocytic cycles and therefore a greater range of host cells that need to be recognized. Although, the identification of invasion genes confirms expectations and to some extent validates the approach, we note that in all of our comparisons the number of genes with no annotated functions, vastly exceeds those with characterized homologs. This emphasises the potential depth of new biology associated with these uncharacterized genes.

Given the absence of a fossil record, the time to the most recent common ancestor was estimated for pairs of species across the *Plasmodium* phylogeny. Although the method is crude, because fixed rates of amino acid evolution are assumed across the tree, we estimate the mammalian and avian lineage of *Plasmodium* split in the order of 10 million years ago, long after the mammals and birds diverged. Combined with new data from the *Laverania* subgenus (Otto et al. 2017), we therefore believe that the idea that the species split of the *Plasmodium* species coincides with their distinctive hosts is no longer tenable.

## Methods

### Collection of parasites and preparation of genomic DNA from *P. gallinaceum* strain 8A

The Institutional Animal Care and Use Committee (IACUC) of the University of California San Diego (UCSD) approved the animal protocol for the production of blood stages of *P. gallinaceum*. The 8A strain (catalogue number MRA-310, ATCC, American Type Culture Collection, Manassas, VA, USA) used in these experiments was originally isolated in 1936 from chickens in Sri Lanka (Brumpt 1937) and has been since kept in laboratories across the world (largely through intraperitoneal passage between chickens and with occasional transmission via infected mosquitoes) as a model species for malaria research in the laboratory (Williams 2005). The *P. gallinaceum* 8A strain was cycled through White Leghorn chickens and *Aedes aegypti* mosquitoes, passage one (P1) parasites (10^5^parasites/chick) were used to infect twenty chickens, and blood was collected as previously described (Patra and Vinetz 2012). Approximately, a total of 100 ml of infected blood sample (>10% parasitemia) was collected, centrifuged, the buffy coat was removed, and the RBC pellet was washed four times with cold phosphate buffered saline (PBS) pH 7.40. Washed RBCs were lysed by saponin (0.05% in PBS) and genomic DNA (gDNA) was extracted using a standard phenol-chloroform method. Because chicken RBC are nucleated only a small proportion of isolated DNA was that of *P. gallinaceum*. Hence Hoechst 33258-Cesium chloride (Cs-Cl) ultracentrifugation was used to separate AT rich *Plasmodium* DNA from the chicken DNA (Dame and Mccutchan 1987). Isolated *P. gallinaceum* DNA was extensively dialyzed against autoclaved Milli-Q water, precipitated with isopropanol and the DNA pellet washed with 70% ethanol. The DNA pellet was suspended in TE (10 mM Tris-HCl, 1 mM EDTA, pH 8) buffer and visualized in 0.7% agarose gel electrophoresis to confirm the quality of the DNA preparation. The DNA was stored at −800C or on dry ice prior to use.

### Host DNA depletion and whole genome sequencing of *P. gallinaceum*

Purified *P. gallinaceum* genomic DNA from a batch prepared in 2003 was used to produce an amplification-free Illumina library of 400-600 base pairs (bp) (Quail et al. 2012) and 100bp paired end reads were generated on an Illumina HiSeq 2000 according to the manufacturer’s standard sequencing protocol. To reduce host contamination and enrich for *P. gallinaceum* DNA, 2 μg of the DNA sample was mixed with 320 μl of methyl binding domain-Fc protein A beads complex (Feehery et al. 2013). The mixture was incubated at room temperature for 15 minutes with gentle rotation. Incubated mixture was placed on a magnetic rack for 3 minutes to separate the beads and the supernatant. A clear supernatant containing enriched *P. gallinaceum* DNA was pipetted in to a clean tube without disturbing the beads. The supernatant was purified using 1.8x volume of Agencourt AMPure^®^ XP beads (Beckman Coulter #A63880) following manufacturer’s instructions. The DNA was eluted in 80 μl of 1x TE buffer (pH 7.5).

An amplification-free Illumina library of 400-600bp was prepared from the enriched genomic DNA (Quail et al. 2012) and 150bp paired end reads were generated on an Illumina MiSeq using v2 chemistry according to the manufacturer’s standard sequencing protocol.

From 20ng of the enriched genomic DNA whole genome amplification (WGA) was performed with REPLI-g Mini Kit (Qiagen) following a modified protocol (Oyola et al. 2014) (Supplemental Information).

This material was then used to prepare a 3-4kb Illumina mate-paired library using an improved (Sanger) mate-paired protocol (Park et al. 2013) and 100bp paired end reads were generated on an Illumina HiSeq 2500 according to the manufacturer’s standard sequencing protocol.

### Collection of parasites, preparation of genomic DNA from *P. relictum* and sequencing

Experimental procedures were approved by the Ethical Committee for Animal Experimentation established by the CNRS under the auspices of the French Ministry of Education and Research (permit number CEEA-LR-1051). *Plasmodium relictum* (lineage SGS1-like, recently renamed DONANA05 (Bensch et al. 2009), GenBank KJ579152) was originally isolated by G. Sorci from wild sparrows (*Passer domesticus*) caught in 2009 in the region of Dijon (France) and subsequently passaged to naïve canaries (*Serinus canaria*) by intra peritoneal injection. The strain was maintained in an animal house by carrying out regular passages between our stock canaries, and occasionally through *Cx pipiens* mosquitoes every ca. 3 weeks (for details see Pigeault et al. 2015).

Midguts were obtained from heavily infected mosquitos (see Supplemental Information).

An amplification-free Illumina library of 400-600 base pair (bp) was prepared from the genomic DNA of infected mosquito midguts and 150bp paired end reads were generated on an Illumina MiSeq using v2 chemistry according to the manufacturer’s standard sequencing protocol.

### Genome assembly and annotation of *P. gallinaceum* and *P. relictum*

Due to the better ratio of parasite versus host, the *P. relictum* assembly generated better contig results. Low quality regions for the reads were clipped with SGA version 0.9.1 (Simpson and Durbin 2012) and contigs were scaffolded with SSPACE (Boetzer et al. 2011). The assembly was improved using PAGIT (Swain et al. 2012) (see Supplemental Information).

The *P. gallinaceum* data were similarly assembled but with more iterative steps of PAGIT, SSPACE and REAPR (made possible due to a 3kb mate-pair library).

Annotation was performed using the Artemis and ACT software (Carver et al. 2013a). Gene model structures were corrected based on orthology and transcriptome data (Lauron et al. 2014). The RNA-seq reads from Lauron et al. were mapped with tophat2 (Kim et al. 2013) against the new *P. gallinaceum* genome. Based on aligned RNA-seq data, 326 gene models were modified and a further eight identified. Functional descriptions were extracted from literature or based on assessment of BLAST and FASTA similarity searches against public databases and searches in protein domain databases such as InterPro and Pfam (Finn et al. 2014). Transmembrane domains were identified using TMHMMv2.0 (Krogh et al. 2001) and Rfamscan (Nawrocki et al. 2015) was used to identify non-coding RNA genes. OrthoMCL 38 (Li et al. 2003) was used to identify orthologs and paralogs.

### Phylogenetic analysis

OrthoMCL v2.0 (default parameters and an inflation parameter of 1.5) was used to identify a total of 881 clusters proteins that were single-copy and present in 19 species of *Apicomplexan* parasites (Supplemental Methods).

For phylogentic analysis (Supplemental Methods), eight independent MCMC chains, each with at least 60,000 steps were run. The final 1500 trees from each chain were concatenated for inference (discarding approximately 20,000 steps per chain as burn-in). To generate the RBP and TE trees, we trimmed the alignments with Gblocks in Seaview version 4.3.1 (Galtier et al. 1996) allowing the loosest settings.

### Dating

G-PhoCS (Gronau et al. 2011) (a Bayesian coalescence method) has been used previously to estimate divergence times of *P. malariae* and *P. malariae-like* (Rutledge et al. 2017).With only a single representative sample for each avian-infective species, it is not possible to use G-PhoCS. We used a method based on a Total Least Squares regression and the existence of a molecular clock specific to *Plasmodium* (Silva et al. 2015) to estimate dating (see Supplemental Methods).

### Transposon analysis

LTRharvest (from GenomeTools v1.5.2) (Ellinghaus et al. 2008) was used to search for putative LTR retrotransposon insertions in the sequence scaffold on which the ORF (4455 bp) in question was located. It successfully identified two flanking LTR sequences of 459 bp (5’LTR) and 469 bp (3’LTR) length and 90% similarity. Subsequent annotation of this element using LTRdigest (Steinbiss et al. 2009) revealed the presence of several profile HMM matches to retrotransposon-associated domains (Gag, protease, reverse transcriptase, RNase H, integrase). Profiles used in this search were collected from the Pfam (Finn et al. 2014) (PF00075, PF00077, PF00078, PF00098, PF00385, PF00552, PF00665, PF00692, PF01021, PF01393, PF02022, PF03732, PF04094, PF04195, PF05380, PF06815, PF06817, PF07253, PF07727, PF08284) and GyDB databases (Llorens et al. 2011). The LTRdigest run also detected a primer binding site of length 15, complementary to a tRNASer (anticodon GCT). For this purpose *P. gallinaceum* tRNA sequences were predicted ab initio using ARAGORN v1.2.36 (Laslett and Canback 2004). Moreover a polypurine tract of length 27 (AAAAAAAAAAAAAAAAAAAAAAAAAGA) was identified manually by examination of the area upstream of the 3’LTR. Filtering and manual inspection of the results of genome-wide LTRharvest/LTRdigest runs discovered at least four more potential near-full-length copies. However, none of these retains a complete ORF.

RepeatMasker (version open-4.0.2, with ABBlast/WUBlast 2.0MP-WashU,-nolow, default sensitivity) was used to identify fragmented insertions of the element in the genome DNA sequence using the DNA sequence of the full-length element as a custom library. All hits of length less than 400 bp were disregarded.

### Prediction of exported proteins

All genes of the reference genomes were analysed for the presence of a PEXEL-motif using the updated HMM algorithm ExportPred v2.0 (Boddey et al. 2013). As a cutoff value 1.5 was used as in (Boddey et al. 2013). To compare genes with PEXEL-motifs between the species we used only orthologous genes with a one-to-one relationship in the 11 reference species.

### Analysis of conserved motifs

To predict new motifs we used MEME version 4.9.1. For the STP1 and *SURFIN* analysis we searched for 96 motifs of the length between 10-150 amino acids on all the existing STP1 and SURFIN sequences of the used 9 genomes. Next, the conceptual proteomes of 9 *Plasmodium* spp were searched for the presence of those STP1/SURFIN meme motifs using fimo, a tool from the meme suit that finds predicted MEME motifs in new sequences (cut-off 1.0E-6; seg used to exclude low complexity amino acid regions). Genes with less than 5 hits were excluded. The output was parsed with a PERL script into a matrix and visualized in R, using the heatmap.2 function and the ward clustering. The phylogenetic trees in Fig. 6 were built with PhyML (Guindon et al. 2009). The alignments for those trees are based on the three meme motifs each. We tried to maximize the occurrence of number of sequences and species for the tree.

For the RBP analysis we took 15 RBP from each species. We chose 15 to have the same number of sequences per species. We joined the two *Laverania* samples and down sampled randomly the amount of sequences if needed. Motifs were predicted with the parameters-nmotifs 96-minw 10-maxw 150.

All structural predictions were performed on the I-Tasser web server (Yang et al. 2015) using default parameter. To determine Pfam domain enrichment we ran InterProScan (Mitchell et al. 2015), and parsed the output in a table for further analysis.

### *d*_N_*/d*_S_ analysis

From the orthologues we aligned the nucleotides after trimming low complexity, with muscle. Amino acids alignments were trimmed with GBlocks (default settings) and against filtered for low complexity with the Seg program. To calculate the *d_N_/d_S_* we used paml and the “Nei & Gojobori” method (Nei and Gojobori 1986). We calculated the *d_N_/d_S_* twice, once just with alignments between the two avian species and once with all eight genomes.

The *d_N_* values were generated in the *d_N_/d_S_* analysis and parsed for each comparison. Intersections between the top 250 genes for each of the six comparisons were visualized with the R package UpSetR (Lex et al. 2014). The product descriptions were obtained from the June 2017 version of *P. falciparum* from GeneDB (Logan-Klumpler et al. 2012).

The GO enrichment with TopGO (Alexa and Rahnenfuhrer 2016) was performed for 17 comparisons and the resulting P-values were multiplied by 17 to adjust for multiple testing.

### Determining the number of RBP copies in *P. relictum*

The long coding sequences (> 7.5 kb) with large blocks of highly similar sequence between RBPs confounded the assembly process and made determining the true number of RBPs a challenge.

In the *P. relictum* assembly, 18 full-length RBP genes were annotated, of which five were pseudogenes. In addition, the *P. relictum* assembly contained numerous gene fragments that were truncated by the assembly process. Despite the high degree of polymorphism, the order of meme motifs in the full length RBP genes was conserved and this information was used to classify and count RBP fragments corresponding to the N and C termini. Of fifteen incomplete genes 8 and 6 were classified as unambiguously representing N-and C-terminal fragments, respectively. To identify copy number variants (CNVs) amongst near-identical RBPs, Illumina reads were mapped back against the genome using BWA (default parameters) (Li and Durbin 2010) and Bamview in Artemis (Carver et al. 2013b). A CNV was assumed where read depth increased by a factor of 2, 3 or 4, relative to the median coverage depth for the whole genome. The occurrence of heterozygous SNPs within a CNV provided additional supporting evidence. Four almost base-perfect copies of one full length RBP had been collapsed into a single copy in the assembly process.

### Expression of *P. gallinaceum* transposase in *P. berghei*

A 6.4 kb fragment carrying the *P. gallinaceum* gag-pol (without the LTR repeats) was synthesised by GeneArt Gene Synthesis service (Thermo Fischer Scientific). The *P. gallinaceum* gag-pol was sub-cloned into pL1694 (obtained from the Leiden Malaria Group) using *BamHI* and *NotI*, placing its expression under control of the constitutive *pbhsp70* promoter. In the resulting vector, the expression cassette *pbhsp70 5’utr:pgtransp:pbhsp70 3’utr* is flanked by homology arms to the *P. berghei* p230p gene. The vector was linearised by *Sac*II prior to transfection into the 1596 *P. berghei* GIMO (<u>g</u>ene <u>i</u>n <u>m</u>arker <u>o</u>ut) mother line (Leiden Malaria Group), with transfectant parasites injected intravenously into Balb/c mice followed by administration of 5-fluorocytosine in the drinking water (Janse et al. 2006; Lin et al. 2011; Braks et al. 2006).

Parasites typically appeared 10 day post-transfection, and were genotyped for integration of the *pbhsp70 5’utr:pgtransp:pbhsp70 3’utr* expression cassette (L1694-GT-F AGCGATAAAAATGATAAACCA, L1694-Pgtransp.-GT-R CGATTGACGCTAAATCATTCGG). Transfection was attempted at four independent occasions, in the presence of a positive control without integration of the *P. gallinaceum* transposase expression cassette ever being detected.

### Data Access

All data generated in this study have been submitted to the European Nucleotide Archive (ENA; http://www.ebi.ac.uk/ena) under accession numbers PRJEB9073 (*P. gallinaceum* 8A genome assembly), PRJEB9074 (*P. relictum* SGS1 genome assembly), PRJEB2470 (*P. gallinaceum* Illumina reads) and PRJEB2579 (*P. relictum* SGS1 Illumina reads). The assembly and annotation are available in GeneDB (http://www.genedb.org) and PlasmoDB (http://www.plasmodb.org).

## Acknowledgments

This work was supported by the Wellcome Trust (grant number 206194). SS and CN were funded by the Wellcome Trust (grant numbers WT099198MA and 104792/Z/14/Z, respectively). We would like to thank Manuel Llinas and Lindsey Orchard for their work on attempting to identify the binding site for the novel avian malaria specific AP2 protein. We would like to thank Joana Silva for her help in dating speciation events.

## References

Alexa A, Rahnenfuhrer J. 2016. topGO: Enrichment Analysis for Gene Ontology. R Package Version 2280.

Asghar M, Hasselquist D, Hansson B, Zehtindjiev P, Westerdahl H, Bensch S. 2015. Chronic infection. Hidden costs of infection: chronic malaria accelerates telomere degradation and senescence in wild birds. Science 347: 436–438.

Atkinson CT, Dusek RJ, Woods KL, Iko WM. 2000. Pathogenicity of avian malaria in experimentally-infected Hawaii Amakihi. J Wildl Dis 36: 197–204.

Bailey TL, Boden M, Buske FA, Frith M, Grant CE, Clementi L, Ren J, Li WW, Noble WS. 2009. MEME SUITE: tools for motif discovery and searching. Nucleic Acids Res 37: W202–208.

Balu B, Chauhan C, Maher SP, Shoue DA, Kissinger JC, Fraser MJ, Adams JH. 2009. piggyBac is an effective tool for functional analysis of the Plasmodium falciparum genome. BMC Microbiol 9: 83.

Bensch S, Canbäck B, DeBarry JD, Johansson T, Hellgren O, Kissinger JC, Palinauskas V, Videvall E, Valkiūnas G. 2016. The Genome of Haemoproteus tartakovskyi and Its Relationship to Human Malaria Parasites. Genome Biol Evol 8: 1361–1373.

Bensch S, Hellgren O, Pérez-Tris J. 2009. MalAvi: a public database of malaria parasites and related haemosporidians in avian hosts based on mitochondrial cytochrome b lineages. Mol Ecol Resour 9: 1353–1358.

Blanquart S, Gascuel O. 2011. Mitochondrial genes support a common origin of rodent malaria parasites and Plasmodium falciparum’s relatives infecting great apes. BMC Evol Biol 11: 70.

Boddey JA, Carvalho TG, Hodder AN, Sargeant TJ, Sleebs BE, Marapana D, Lopaticki S, Nebl T, Cowman AF. 2013. Role of plasmepsin V in export of diverse protein families from the Plasmodium falciparum exportome. Traffic Cph Den 14: 532–550.

Boetzer M, Henkel CV, Jansen HJ, Butler D, Pirovano W. 2011. Scaffolding pre-assembled contigs using SSPACE. Bioinforma Oxf Engl 27: 578–579.

Borner J, Pick C, Thiede J, Kolawole OM, Kingsley MT, Schulze J, Cottontail VM, Wellinghausen N, Schmidt-Chanasit J, Bruchhaus I, et al. 2016. Phylogeny of haemosporidian blood parasites revealed by a multi-gene approach. Mol Phylogenet Evol 94: 221–231.

Braks JAM, Franke-Fayard B, Kroeze H, Janse CJ, Waters AP. 2006. Development and application of a positive-negative selectable marker system for use in reverse genetics in Plasmodium. Nucleic Acids Res 34: e39.

Brumpt É. 1937. Schizogonie parfois intense du Plasmodium gallinaceum dans les cellules endotheliales des poules. C R Soc Biol Paris 810–813.

Carver T, Harris SR, Otto TD, Berriman M, Parkhill J, McQuillan JA. 2013a. BamView: visualizing and interpretation of next-generation sequencing read alignments. Brief Bioinform 14: 203–212.

Carver T, Harris SR, Otto TD, Berriman M, Parkhill J, McQuillan JA. 2013b. BamView: visualizing and interpretation of next-generation sequencing read alignments. Brief Bioinform 14: 203–212.

Cunningham D, Lawton J, Jarra W, Preiser P, Langhorne J. 2010. The pir multigene family of Plasmodium: antigenic variation and beyond. Mol Biochem Parasitol 170: 65–73.

Dame JB, Mccutchan TF. 1987. Plasmodium falciparum: Hoechst Dye 33258-CsCl ultracentrifugation for separating parasite and host DNAs. Exp Parasitol 64: 264–266.

Ellinghaus D, Kurtz S, Willhoeft U. 2008. LTRharvest, an efficient and flexible software for de novo detection of LTR retrotransposons. BMC Bioinformatics 9: 18.

Feehery GR, Yigit E, Oyola SO, Langhorst BW, Schmidt VT, Stewart FJ, Dimalanta ET, Amaral-Zettler LA, Davis T, Quail MA, et al. 2013. A method for selectively enriching microbial DNA from contaminating vertebrate host DNA. PloS One 8: e76096.

Fell VL, Schild-Poulter C. 2015. The Ku heterodimer: function in DNA repair and beyond. Mutat Res Rev Mutat Res 763: 15–29.

Fernandez-Becerra C, Yamamoto MM, Vêncio RZN, Lacerda M, Rosanas-Urgell A, del Portillo HA. 2009. Plasmodium vivax and the importance of the subtelomeric multigene vir superfamily. Trends Parasitol 25: 44–51.

Finn RD, Bateman A, Clements J, Coggill P, Eberhardt RY, Eddy SR, Heger A, Hetherington K, Holm L, Mistry J, et al. 2014. Pfam: the protein families database. Nucleic Acids Res 42: D222–230.

Galtier N, Gouy M, Gautier C. 1996. SEAVIEW and PHYLO_WIN: two graphic tools for sequence alignment and molecular phylogeny. Comput Appl Biosci CABIOS 12: 543–548.

Gardner MJ, Hall N, Fung E, White O, Berriman M, Hyman RW, Carlton JM, Pain A, Nelson KE, Bowman S, et al. 2002. Genome sequence of the human malaria parasite Plasmodium falciparum. Nature 419: 498–511.

Garnham PCC. 1966. Malaria parasites and other haemosporidia. Blackwell Scientific Publishers, Oxford, UK.

Glaizot O, Fumagalli L, Iritano K, Lalubin F, Van Rooyen J, Christe P. 2012. High prevalence and lineage diversity of avian malaria in wild populations of great tits (Parus major) and mosquitoes (Culex pipiens). PloS One 7: e34964.

Goel S, Palmkvist M, Moll K, Joannin N, Lara P, Akhouri RR, Moradi N, öjemalm K, Westman M, Angeletti D, et al. 2015. RIFINs are adhesins implicated in severe Plasmodium falciparum malaria. Nat Med 21: 314–317.

Gronau I, Hubisz MJ, Gulko B, Danko CG, Siepel A. 2011. Bayesian inference of ancient human demography from individual genome sequences. Nat Genet 43: 1031–1034.

Guindon S, Delsuc F, Dufayard J-F, Gascuel O. 2009. Estimating maximum likelihood phylogenies with PhyML. Methods Mol Biol Clifton NJ 537: 113–137.

Haltiwanger BM, Matsumoto Y, Nicolas E, Dianov GL, Bohr VA, Taraschi TF. 2000. DNA base excision repair in human malaria parasites is predominantly by a long-patch pathway. Biochemistry (Mosc) 39: 763–772.

Huff CG. 1969. Exoerythrocytic stages of avian and reptilian malarial parasites. Exp Parasitol 24: 383–421.

Huff CG, Bloom W. 1935. A malarial parasite infecting all blood and blood-forming cells in birds. J Infect Dis 57: 315–336.

Janse CJ, Ramesar J, Waters AP. 2006. High-efficiency transfection and drug selection of genetically transformed blood stages of the rodent malaria parasite Plasmodium berghei. Nat Protoc 1: 346–356.

Janssen CS, Phillips RS, Turner CMR, Barrett MP. 2004. Plasmodium interspersed repeats: the major multigene superfamily of malaria parasites. Nucleic Acids Res 32: 5712–5720.

Kim D, Pertea G, Trapnell C, Pimentel H, Kelley R, Salzberg SL. 2013. TopHat2: accurate alignment of transcriptomes in the presence of insertions, deletions and gene fusions. Genome Biol 14: R36.

Krogh A, Larsson B, von Heijne G, Sonnhammer EL. 2001. Predicting transmembrane protein topology with a hidden Markov model: application to complete genomes. J Mol Biol 305: 567–580.

Kumar S, Hedges SB. 1998. A molecular timescale for vertebrate evolution. Nature 392: 917–920.

Lachish S, Knowles SCL, Alves R, Wood MJ, Sheldon BC. 2011. Fitness effects of endemic malaria infections in a wild bird population: the importance of ecological structure. J Anim Ecol 80: 1196–1206.

Lapointe DA, Atkinson CT, Samuel MD. 2012. Ecology and conservation biology of avian malaria. Ann N Y Acad Sci 1249: 211–226.

Laslett D, Canback B. 2004. ARAGORN, a program to detect tRNA genes and tmRNA genes in nucleotide sequences. Nucleic Acids Res 32: 11–16.

Lauron EJ, Aw Yeang HX, Taffner SM, Sehgal RNM. 2015. De novo assembly and transcriptome analysis of Plasmodium gallinaceum identifies the Rh5 interacting protein (ripr), and reveals a lack of EBL and RH gene family diversification. Malar J 14: 296.

Lauron EJ, Oakgrove KS, Tell LA, Biskar K, Roy SW, Sehgal RN. 2014. Transcriptome sequencing and analysis of Plasmodium gallinaceum reveals polymorphisms and selection on the apical membrane antigen-1. Malar J 13: 382.

Levin II, Zwiers P, Deem SL, Geest EA, Higashiguchi JM, Iezhova TA, Jiménez-Uzcátegui G, Kim DH, Morton JP, Perlut NG, et al. 2013. Multiple lineages of Avian malaria parasites (Plasmodium) in the Galapagos Islands and evidence for arrival via migratory birds. Conserv Biol J Soc Conserv Biol 27: 1366–1377.

Lex A, Gehlenborg N, Strobelt H, Vuillemot R, Pfister H. 2014. UpSet: Visualization of Intersecting Sets. IEEE Trans Vis Comput Graph 20: 1983–1992.

Li H, Durbin R. 2010. Fast and accurate long-read alignment with Burrows-Wheeler transform. Bioinforma Oxf Engl 26: 589–595.

Li L, Stoeckert CJ, Roos DS. 2003. OrthoMCL: identification of ortholog groups for eukaryotic genomes. Genome Res 13: 2178–2189.

Lin J, Annoura T, Sajid M, Chevalley-Maurel S, Ramesar J, Klop O, Franke-Fayard BMD, Janse CJ, Khan SM. 2011. A novel “gene insertion/marker out” (GIMO) method for transgene expression and gene complementation in rodent malaria parasites. PloS One 6: e29289.

Ling K-H, Rajandream M-A, Rivailler P, Ivens A, Yap S-J, Madeira AMBN, Mungall K, Billington K, Yee W-Y, Bankier AT, et al. 2007. Sequencing and analysis of chromosome 1 of Eimeria tenella reveals a unique segmental organization. Genome Res 17: 311–319.

Llorens C, Futami R, Covelli L, DomÍnguez-Escribá L, Viu JM, Tamarit D, Aguilar-RodrÍguez J, Vicente-Ripolles M, Fuster G, Bernet GP, et al. 2011. The Gypsy Database (GyDB) of mobile genetic elements: release 2.0. Nucleic Acids Res 39: D70–74.

Logan-Klumpler FJ, De Silva N, Boehme U, Rogers MB, Velarde G, McQuillan JA, Carver T, Aslett M, Olsen C, Subramanian S, et al. 2012. GeneDB—an annotation database for pathogens. Nucleic Acids Res 40: D98–108.

Mancio-Silva L, Slavic K, Grilo Ruivo MT, Grosso AR, Modrzynska KK, Vera IM, Sales-Dias J, Gomes AR, MacPherson CR, Crozet P, et al. 2017. Nutrient sensing modulates malaria parasite virulence. Nature 547: 213–216.

Marshall EK. 1942. Chemotherapy of avian malaria. Physiol Rev 22: 190–204.

Marti M, Good RT, Rug M, Knuepfer E, Cowman AF. 2004. Targeting malaria virulence and remodeling proteins to the host erythrocyte. Science 306: 1930–1933.

Merino EF, Fernandez-Becerra C, Durham AM, Ferreira JE, Tumilasci VF, d’Arc-Neves J, da Silva-Nunes M, Ferreira MU, Wickramarachchi T, Udagama-Randeniya P, et al. 2006. Multicharacter population study of the vir subtelomeric multigene superfamily of Plasmodium vivax, a major human malaria parasite. Mol Biochem Parasitol 149: 10–16.

Mitchell A, Chang H-Y, Daugherty L, Fraser M, Hunter S, Lopez R, McAnulla C, McMenamin C, Nuka G, Pesseat S, et al. 2015. The InterPro protein families database: the classification resource after 15 years. Nucleic Acids Res 43: D213–221.

Nawrocki EP, Burge SW, Bateman A, Daub J, Eberhardt RY, Eddy SR, Floden EW, Gardner PP, Jones TA, Tate J, et al. 2015. Rfam 12.0: updates to the RNA families database. Nucleic Acids Res 43: D130–137.

Neafsey DE, Galinsky K, Jiang RHY, Young L, Sykes SM, Saif S, Gujja S, Goldberg JM, Young S, Zeng Q, et al. 2012. The malaria parasite Plasmodium vivax exhibits greater genetic diversity than Plasmodium falciparum. Nat Genet 44: 1046–1050.

Nei M, Gojobori T. 1986. Simple methods for estimating the numbers of synonymous and nonsynonymous nucleotide substitutions. Mol Biol Evol 3: 418–426.

Niang M, Bei AK, Madnani KG, Pelly S, Dankwa S, Kanjee U, Gunalan K, Amaladoss A, Yeo KP, Bob NS, et al. 2014. STEVOR is a Plasmodium falciparum erythrocyte binding protein that mediates merozoite invasion and rosetting. Cell Host Microbe 16: 81–93.

Otto TD, Dillon GP, Degrave WS, Berriman M. 2011. RATT: Rapid Annotation Transfer Tool. Nucleic Acids Res 39: e57.

Otto TD, Gilabert A, Crellen T, Böhme U, Arnathau C, Sanders M, Oyola S, Okauga AP, Boundenga L, Wuillaume E, et al. 2017. Genomes of an entire Plasmodium subgenus reveal paths to virulent human malaria. bioRxiv doi: https://doi.org/10.1101/095679.

Otto TD, Rayner JC, Böhme U, Pain A, Spottiswoode N, Sanders M, Quail M, Ollomo B, Renaud F, Thomas AW, et al. 2014. Genome sequencing of chimpanzee malaria parasites reveals possible pathways of adaptation to human hosts. Nat Commun 5: 4754.

Oyola SO, Gu Y, Manske M, Otto TD, O’Brien J, Alcock D, Macinnis B, Berriman M, Newbold CI, Kwiatkowski DP, et al. 2013. Efficient depletion of host DNA contamination in malaria clinical sequencing. J Clin Microbiol 51: 745–751.

Oyola SO, Manske M, Campino S, Claessens A, Hamilton WL, Kekre M, Drury E, Mead D, Gu Y, Miles A, et al. 2014. Optimized whole-genome amplification strategy for extremely AT-biased template. DNA Res IntJ Rapid Publ Rep Genes Genomes 21: 661–671.

Pain A, Böhme U, Berry AE, Mungall K, Finn RD, Jackson AP, Mourier T, Mistry J, Pasini EM, Aslett MA, et al. 2008. The genome of the simian and human malaria parasite Plasmodium knowlesi. Nature 455: 799–803.

Park N, Shirley L, Gu Y, Keane TM, Swerdlow H, Quail MA. 2013. An improved approach to mate-paired library preparation for Illumina sequencing. Methods Gener Seq 1.

Patra KP, Vinetz JM. 2012. New Ultrastructural Analysis of the Invasive Apparatus of the Plasmodium Ookinete. Am J Trop Med Hyg 87: 412–417.

Perkins SL. 2014. Malaria’s many mates: past, present, and future of the systematics of the order Haemosporida. J Parasitol 100: 11–25.

Perkins SL, Schaer J. 2016. A Modern Menagerie of Mammalian Malaria. Trends Parasitol 32: 772–782.

Pick C, Ebersberger I, Spielmann T, Bruchhaus I, Burmester T. 2011. Phylogenomic analyses of malaria parasites and evolution of their exported proteins. BMC Evol Biol 11: 167.

Pigeault R, Nicot A, Gandon S, Rivero A. 2015. Mosquito age and avian malaria infection. Malar J 14: 383.

Quail MA, Smith M, Coupland P, Otto TD, Harris SR, Connor TR, Bertoni A, Swerdlow HP, Gu Y. 2012. A tale of three next generation sequencing platforms: comparison of Ion Torrent, Pacific Biosciences and Illumina MiSeq sequencers. BMC Genomics 13: 341.

Raffaela G, Marchiafava E. 1944. Avian malaria: a new lease of life for an old experimental model to study the evolutionary ecology of Plasmodium. Ann Soc Belg Med Trop 323–330.

Reid AJ, Blake DP, Ansari HR, Billington K, Browne HP, Bryant JM, Dunn M, Hung SS, Kawahara F, Miranda-Saavedra D, et al. 2014. Genomic analysis of the causative agents of coccidiosis in domestic chickens. Genome Res.

Rutledge GG, Böhme U, Sanders M, Reid AJ, Cotton JA, Maiga-Ascofare O, Djimdé AA, Apinjoh TO, Amenga-Etego L, Manske M, et al. 2017. Plasmodium malariae and P. ovale genomes provide insights into malaria parasite evolution. Nature 542: 101–104.

Saito F, Hirayasu K, Satoh T, Wang CW, Lusingu J, Arimori T, Shida K, Palacpac NMQ, Itagaki S, Iwanaga S, et al. 2017. Immune evasion of Plasmodium falciparum by RIFIN via inhibitory receptors. Nature 552: 101–105.

Silva JC, Egan A, Arze C, Spouge JL, Harris DG. 2015. A New Method for Estimating Species Age Supports the Coexistence of Malaria Parasites and Their Mammalian Hosts. Mol Biol Evol 32: 1354–1364.

Simpson JT, Durbin R. 2012. Efficient de novo assembly of large genomes using compressed data structures. Genome Res 22: 549–556.

Spence PJ, Jarra W, Lévy P, Reid AJ, Chappell L, Brugat T, Sanders M, Berriman M, Langhorne J. 2013. Vector transmission regulates immune control of Plasmodium virulence. Nature 498: 228–231.

Springer WT. 1996. Other blood and tissue protozoa. In Calnek, B.W, Beard, C.W., McDougald, L.R., Saif, Y.M. (Eds.). Diseases of Poultry, pp. 900–911, Ames, IA: Iowa State University Press.

Steinbiss S, Willhoeft U, Gremme G, Kurtz S. 2009. Fine-grained annotation and classification of de novo predicted LTR retrotransposons. Nucleic Acids Res 37: 7002–7013.

Swain MT, Tsai IJ, Assefa SA, Newbold C, Berriman M, Otto TD. 2012. A post-assembly genome-improvement toolkit (PAGIT) to obtain annotated genomes from contigs. Nat Protoc 7: 1260–1284.

Templeton TJ, Enomoto S, Chen W-J, Huang C-G, Lancto CA, Abrahamsen MS, Zhu G. 2010. A genome-sequence survey for Ascogregarina taiwanensis supports evolutionary affiliation but metabolic diversity between a Gregarine and Cryptosporidium. Mol Biol Evol 27: 235–248.

Valkiunas G. 2004. Avian Malaria Parasites and other Haemosporidia. CRC Press.

Ward JH. 1963. Hierarchical Grouping to Optimize an Objective Function. J Am Stat Assoc 58: 236–244.

Waters AP, Higgins DG, McCutchan TF. 1991. Plasmodium falciparum appears to have arisen as a result of lateral transfer between avian and human hosts. Proc Natl Acad Sci USA 88: 3140–3144.

Williams RB. 2005. Avian malaria: clinical and chemical pathology of Plasmodium gallinaceum in the domesticated fowl Gallus gallus. Avian Pathol J WVPA 34: 29–47.

Winter G, Kawai S, Haeggström M, Kaneko O, von Euler A, Kawazu S, Palm D, Fernandez V, Wahlgren M. 2005. SURFIN is a polymorphic antigen expressed on Plasmodium falciparum merozoites and infected erythrocytes. J Exp Med 201: 1853–1863.

Yang J, Yan R, Roy A, Xu D, Poisson J, Zhang Y. 2015. The I-TASSER Suite: protein structure and function prediction. Nat Methods 12: 7–8.

